# Stress priming affects fungal competition – evidence from a combined experimental and modeling study

**DOI:** 10.1101/2020.03.04.976357

**Authors:** Felix Wesener, Aleksandra Szymczak, Matthias C. Rillig, Britta Tietjen

## Abstract

Priming, an inducible stress defense strategy that prepares an organism for an impending stress event, is common in microbes and has been studied mostly in isolated organisms or populations. How the benefits of priming change in the microbial community context and, vice versa, whether priming influences competition between organisms, remains largely unknown. In this combined experimental and modeling study, we developed a cellular automaton model based on dedicated data of different isolates of soil fungi in isolation and pairwise competition experiments. With the model, we simulated growth of the ascomycete *Chaetomium elatum* competing against other fungi to understand which species traits influence the benefit of priming and the effect of priming on competition. We showed that competition changes the priming benefit compared to isolated growth, and that it depends not only on the primeable species itself, but also on the competitors’ traits such as growth rate, primeability and stress susceptibility. In addition, we showed that priming benefits were not always reflected in the competitive outcome. With this study, we transferred insights on priming from studies in isolation to the community context. This is an important step towards understanding the role of inducible defenses in microbial community assembly and composition.

## Introduction

Priming is a stress defense mechanism that enables an organism to remember an environmental cue and to build up an enhanced stress response to a potentially stronger future stress. Primed defense mechanisms have been observed across many microbial taxa (see meta-analysis by Andrade-Linares, Lehmann and Rillig 2016), most of which have focused on the molecular processes that underlie priming. Complementary to research on priming processes, understanding the role of priming in stress ecology is an important step to comprehend how priming might change the effect of stressors on species fitness and community development. At the ecological level, it is still unclear how the ability of an individual to be primed, termed primeability, might influence community development and, vice versa, how the community context affects the benefits of priming.

Microbial priming is a defense strategy found in bacteria (Koutsoumanis and Sofos, 2004; Mitchell *et al.*, 2009; Cebrián *et al.*, 2010; Hernández *et al.*, 2012), archaea (Trent, 1996), as well as fungi (Alvarez-Peral *et al.*, 2002; Berry and Gasch, 2008; Rangel *et al.*, 2008; Mitchell *et al.*, 2009; Guhr *et al.*, 2017). Especially fungi are suitable model organisms to study the effects of priming under different conditions, as many isolates exhibit varying degrees of primeability (Szymczak *et al.*, 2019) and memory length (Diana R. Andrade-Linares *et al.*, 2016). In nature, isolated growth of fungi is rare, usually occurring only when new territory is colonized (Boddy, 2000), and fungi normally live in highly complex communities of different species that compete for space and display a broad range of mostly antagonistic interactions (Boddy, 2000; Toljander *et al.*, 2006; Hiscox and Boddy, 2017), which influence community composition (Boddy, 2000, 2001). Several studies have shown that fungal combative ability is not only dependent on the species that interact, but also on environmental factors such as resource availability (Stahl and Christensen, 1992; Falconer *et al.*, 2008) or temperature (Boddy *et al.*, 1985; Schoeman *et al.*, 1996; Toljander *et al.*, 2006; Hiscox, Clarkson, *et al.*, 2016), and temperature changes can even lead to reversed competitive outcomes (Crowther *et al.*, 2012). Therefore, we expect that heat priming, which affects species of distinct primeability differently in their response to heat stress, has an impact on fungal community development.

Experimental research on priming usually requires time-intensive multifactorial setups, in which organisms experience, apart from control conditions, a stress with and without preceding priming cue, as well as a priming cue without subsequent stress (Hilker *et al.*, 2016). Here, simulation models can complement laboratory experiments by testing different environmental factors and species traits, e.g. imitating conditions or species combinations that could not be investigated empirically. A modeling approach thus allows a systematic investigation of distinct costs and benefits of priming for an organism. Using a mathematical model of microbes in competition, Rillig *et al.* (2015) could show that priming is beneficial more often under community conditions compared to species investigated under isolation. A follow-up study (Wesener and Tietjen, 2019), additionally showed that different strategies to reach an enhanced stress response are of different benefit. Especially the stress duration determined if an early or fast buildup of the response was most beneficial or a stronger response. However, a general understanding of how the benefit of priming can change under competition and how priming influences community structure, is still missing. To fill this gap, we carried out an experiment to collect dedicated data and developed a cellular automaton model simulating the growth of fungal colonies in isolation and in pairwise interactions. Our model is based on experimental data of the ascomycete *Chaetomium elatum* as focal species and various competitors of *C. elatum*, experiencing a mild temperature stimulus and/or heat stress. It can successfully reproduce the growth dynamics of two competing soil fungi under primed or non-primed conditions. To gather a general understanding of priming impacts on fungal communities, we systematically varied different traits of the species competing with *C. elatum* and observed how the priming benefit of *C. elatum* changed depending on its competitor. The specific aims of the study are i) to identify fungal traits that affect the pay-off of priming by comparing the benefit of *C. elatum* in dual cultures with various competitors and ii) to assess the influence of priming on competitive success.

## Methods

We carried out an experiment on six soil fungal species growing in isolation and in dual-species settings to determine the intrinsic growth rate of each species measured as colony diameter extension. For these settings, we determined how the growth of species is altered by heat stress and by priming towards this stress. We used parts of the data (growth under different stress treatments in isolation and unstressed growth under competition) to parameterize growth rates of the model. We then validated our model by comparing the model output with the experimental data on dual-culture growth under stress and priming. Finally, we used the validated model to systematically assess the effect of fungal species traits and of competition on the benefits of priming. Moreover, the model served to investigate how priming influences competition between species.

### Experimental setup

For the laboratory experiment, six soil fungal species (Ascomycetes *Chaetomium elatum, Fusarium redolens, Fusarium oxysporum* and *Truncatella angustata*, Basidiomycete *Pleurotus sapidus* and Mucoromycete *Morteriella elongata*) were grown in isolation and in competition in a full-factorial design of priming and triggering for heat stress. The soil saprotrophic fungi were originally taken from a grassland site in Mallnow (Mallnow Lebus, Brandenburg, Germany, 52°27.778’ N, 14°29.349’ E). All fungi were grown under constant conditions of 22 °C in a Petri dish of 90 mm diameter on potato dextrose agar (PDA). The durations and intensities of priming stimulus and strong heat stress (triggering stimulus) were determined in a pre-experiment. As priming stimulus, we used a mild, non-detrimental heat stress of 35 °C for two hours, as triggering stimulus, we used a temperature of 45 °C for two hours. After receiving a priming/triggering stimulus, the temperature was set back to 22 °C. The full-factorial combination of priming and triggering stimuli resulted in the following treatments: i) A control treatment (C), simulating constant conditions of 22 °C with no disturbance, ii) a priming-only treatment (P), in which a fungus experienced the priming stimulus after one day of undisturbed growth, iii) a triggering-only treatment (T), in which a triggering heat stress was applied, and iv) a primed stress treatment (PT), in which the priming stimulus was immediately followed by a triggering stimulus.

For the single-species experiments, the colony diameter was measured once per day to determine species-specific colony extension rates. Measurements were taken until the colonies reached the edges of the Petri dish or, for slower growing individuals, for 14 consecutive days.

To investigate fungal growth under competition, plugs of mycelium of *C. elatum* were inoculated pairwise with each of the five other species equally distant from each other and from the border of the Petri dish. As soon as the two colonies touched, the same treatments on priming and triggering as in the single species experiments were applied. In pairwise cultures, the individual colony shapes were not circular. Therefore, instead of diameter measurements the surface area of each colony was scanned. Total colony size was then determined in ImageJ (Schneider *et al.*, 2012) to assess colony extension rates. Scanning took place four times in total, until the Petri dish was filled.

Eight replicates per fungus/competition setup and treatment were measured.

### Determination of colony extension rates

We determined the species-specific growth rates by measuring the change of colony diameter (single species) or colony area (dual cultures) over time in all four experimental treatments. We compared the growth in the C and P treatment to determine whether a priming stimulus in itself would already reduce fungal growth. A significant reduction of growth in the P treatment would imply that priming inflicts costs on a primed species on the level of growth, in case a priming stimulus was not followed by a triggering stress. The comparison between C and T treatment revealed the effect of stress on growth, while comparing PT and T treatment quantified how much better a species grows, if primed before stressed.

For the C and P treatment, a linear fit was applied to the daily diameter values of the single species experiments resulting in colony growth measured as colony diameter change [mm/day]. For the T and PT treatments, we detected the occurrence of a stress-induced lag phase, i.e. a period of no growth, and determined the growth of the following phase. To do so, we compared the colony diameter distribution of the single species replicates on day 1 (day of treatment) with each respective following day using a paired t-test after testing for normality and homoscedasticity. We measured the duration of the lag phase *L*_*si,T*_ and *L*_*si,PT*_ as the time period following a T and PT treatment, in which the colony diameter does not differ significantly from day 1. Growth rates were determined for the post-lag growth phase with a linear fit. The exact time point of transition from lag phase to growth was the intersection of the lag and growth fit lines.

Of the six species that were primed experimentally, four did not show a significant difference in growth between control and priming treatment, i.e. no priming costs occurred on the level of growth (Table 1). The other two species, *F. oxysporum* and *T. angustata*, showed a slight overall increase in growth. To reduce the complexity of our model, we thus chose to exclude priming costs for further analyses. The effects of heat stress were similar across all species but two: all fungi exhibited a lag phase without any growth, and four of the six species did not show a change in growth rate after the lag had ended compared to unstressed growth. Solely *T. angustata* showed growth that was different by more than 10% and even higher than under control conditions, while *F. oxysporum* showed a slight reduction in growth after the lag phase.

**Table 1.** Experimentally measured values of growth rates and their relative changes and lag phases after a stress stimulus. The significance levels between growth rates were assessed by a paired t-test * P ≤ 0.05, ** P ≤ 0.01, relative changes of 1 indicated a non-significant change in growth rate. Abbreviations: C: control treatment without stress, P: primed treatment with a mild stress, T: triggered treatment with a strong stress, PT: primed and triggered treatment.

| Species | Control growth rate $g_{si}$ [mm/d] | Rel. change in growth T/C | Rel. change in growth PT/T | Rel. change in growth P/C | Non-primed lag phase $L_{si,T}$ [d] | Primed lag phase $L_{si,PT}$ [d] |
| --- | --- | --- | --- | --- | --- | --- |
| *C. elatum* | 9.38 | 1 | 1 | 1 | 3.17 | 1.9 |
| *M. elongata* | 14.74 | 1 | 1 | 1 | 2.82 | 2.82 |
| *F. oxysporum* | 10.57 | 0.91 (**) | 1 | 1.046 (*) | 0.55 | 0.12 |
| *F. redolens* | 6.63 | 1 | 1 | 1 | 0.71 | 0.01 |
| *P. sapidus* | 3.58 | 1 | 1 | 1 | 1.93 | 1.56 |
| *T. angustata* | 7.13 | 1.15 (**) | 1 | 1.10 (*) | 1.577 | 1.03 |

When being primed, the lag phase after experiencing stress was shorter in five species or remained equal in *M. elongata*, and the growth rate after the end of the lag phase did not differ. Again, to avoid unnecessary model complexity, we assumed no difference in the growth rate after stress-induced lag phases for both T and PT treatments.

### Simulation Model

To simulate a fungal colony growing in a Petri dish, we developed a cellular automaton model and introduced the experimentally determined growth rates into this model. In the following section, we describe the model and how we converted the measured growth rates of the experiment, which are constant over time and space, to the necessary discrete units of time and space of the cellular automaton.

Our model represents a Petri dish, i.e. a circular area, with an inner diameter *d* = 86.5 *mm* containing one or two fungal colonies. The area of the Petri dish is divided into square grid cells with a side length of *r*_*spat*_ = 0.5 *mm*, leading to 173 grid cells along the diameter of the Petri dish. To mimic the laboratory experiments, the initial colony diameter of a fungus is set to *d* = 6*mm*. Colonies in isolation are placed into the center of the Petri dish. Colonies in pairwise experiments are placed on the horizontal diameter equidistant from each other and the border of the Petri dish, i.e. the distance between the centers of the colonies is about 28.5 mm.

Simulation of cell division and colony growth follows the cellular automaton model of Gerlee and Anderson (2007). To realize radial extension of the initial colonies, each grid cell of the model is assigned one of three states: empty, occupied by an active (growing) fungal cell or occupied by a dormant fungal cell. Fungal cells with empty neighboring grid cells (neighbors being all grid cells in the Moore neighborhood of a respective cell) that could thus still spread are considered active cells, while fungal cells without any empty neighboring grid cells turn dormant. Only active fungal cells conduct cell division, i.e. produced new daughter cells on one of the empty neighboring grid cells, leading to an increase in colony area. Therefore, active fungal cells can solely be found in the periphery of a fungal colony. Each time step, the active fungal cells are updated and behaved accordingly, i.e. active cells that are no longer in the colony circumference change to the dormant state.

The temporal resolution *r*_*temp*_ is one hour. To match measured growth rates, it is necessary to determine the frequency of cell division, for which we introduce a linear increasing maturation value *m(t)* for each active cell. Cell division occurs when a cell reaches a maturation age of *m ≥* 1. The increase in *m*, Δ*m,* is calculated based on the measured growth rate *g*_*si*_ relative to the temporal and spatial resolution:

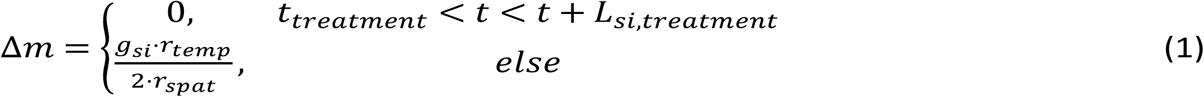

with *si* referring to the simulated species and *treatment* referring to the primed only or primed and triggered treatment. Growth rate is corrected by factor ½, since the measured data of *g*_*si*_ include diameter growth into two directions, while simulated cell division for each side of the colony periphery is calculated separately. In case of heat stress (T or PT treatment), the maturation value remains constant for the duration of the post-stress lag phase, which starts after the application of a heat pulse (at *t*_*T*_) or a priming stimulus followed by a heat pulse (at *t*_*PT*_), and ends after the duration of the lag phase (*L*_*si,T*_ or *L*_*si,PT*_, respectively).

When a fungal cell reaches maturation age of *m* ≥ 1, the cell divides and fills a random empty neighboring grid cell with a higher priority on the immediate four neighboring grid cells, leading to an increase in area. The maturation value is then reduced by 1. The average division number corresponds to the measured radial colony extension. The daughter cell inherits its mother’s new maturation age adjusted by a random variation term with a standard deviation of *σ* = *m/*2. If at the point of division none of the neighboring grid cells is empty, the division failed and the fungal cell changes to the dormant state.

Apart from competition for space, no other forms of interactions are included. The model setup leads to deadlock as only possible competitive outcome, which is a clear separation of space between competing species and the most common competitive outcome of mycelial interactions (Stahl and Christensen, 1992; Schoeman *et al.*, 1996; Hiscox *et al.*, 2018). The cellular automaton model was implemented in NetLogo 6.1.0 (Wilensky, 1999) and analysed using R (R., 2018) and the nlrx package (Salecker *et al.*, 2019).

### Model Parameterization

The following model parameters are based on experimental data of single-species and are specific for each species *si*: i) The intrinsic colony extension rate in isolation *g*_*si*_ [mm/day], ii) thestress susceptibility definedas thelength of the fungistatic lag phase under heat stress *sus*_*si*_ = *L*_*si,T*_, and iii) the primeability of a species describing the reduction of the lag phase if primed before stressed 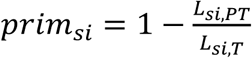. A primeability value of *prim*_*si*_ = 1 describes full primeability, i.e. a reduction of the lag phase to zero, while a primeability of *prim*_*si*_ = 0 applies to non-primeable species that exhibit the same lag phase under T and PT treatment.

Because fungi change their growth rates in dual cultures depending on their competitor (Stahl and Christensen 1992), we adjusted the growth rate *g*_*si,ci*_ of species *si* under competition with competitor *ci* by re-calibrating growth to match the growth observed in the dual culture control experiments. To determine the change in growth rates under competition, we applied a fit to non-stressed conditions, and only on data points before both fungal colonies touched.

With this, we can differentiate the effect that arises due to interactions at distance from the effect of direct competition for space. The latter is already implemented in the model, as simulated fungal cells can only colonize empty grid cells. All parameters are given in Table 2.

**Table 2.** Model parameters and their description.

| Parameter | Description | Source | Unit |
| --- | --- | --- | --- |
| $g_{si}$ | intrinsic colony extension rate in isolation | measured | $mm/d$ |
| $g_{si,ci}$ | intrinsic colony extension rate under competition | measured | $mm/d$ |
| $L_{si,T}$ | duration of a phase of no growth after a 2 h heat stress | measured | d |
| $L_{si,PT}$ | duration of a lag phase of no growth if primed before stressed | measured | d |
| $sus_{si}$ | stress susceptibility | $sus_{si} = L_{si,T}$ | d |
| $prim_{si}$ | stress primeability (reduction of the stress-induced lag phase if primed) | $prim_{si} = 1 - \frac{L_{si,PT}}{L_{si,T}}$ | - |

### Simulation Experiments

To validate the model, we simulated the growth of *C. elatum* as focal species in isolation and under competition with each one of the other five fungi under stress treatment (T) and primed- and-stressed treatment (PT). As model output, we determined the colony area of *C. elatum A_C. elatum_* = *N*_*cells*_ · *r*_*spat*_2 [mm^2^], which serves as a relative measure of fitness. The benefit of priming for *C. elatum* was then described as the colony area of a fungus under stress (T) compared to the area of a primed colony under stress (PT):

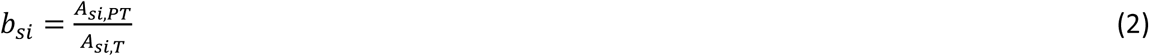

To assess model performance, the colony growth of *C. elatum* with each of its respective competitors as well as the simulated benefit was compared to experimental data.

Subsequently, we performed simulation experiments to determine which species-specific traits influence the benefit of priming and the competition between two species under priming. First, we simulated the growth of *C. elatum* in competition with an artificial species initially exhibiting the same trait values as *C. elatum* until the Petri dish was filled or up to a maximum of 15 days. To examine the effect of specific traits and trait combinations on the priming benefit, we systematically varied the growth rate, stress susceptibility and primeability values of the competitor and measured the benefit *b* of *C. elatum*. Because the growth of *C. elatum* under competition proved to be variable depending on its competitor, we also varied the growth rate of *C. elatum*, while all other trait values of *C. elatum* remained fixed.

Secondly, to determine the effect of priming on a simple community, i.e. which competitor benefits more from priming, we measured the relative benefit

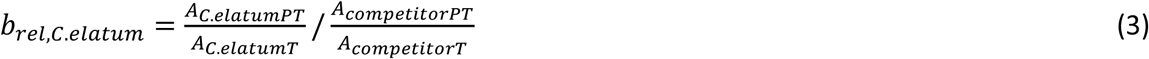

which compares the benefit of both competitors.

Equation (3) can be rearranged and log-transformed to describe the change in colony ratio 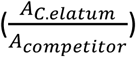 between T and PT treatment:

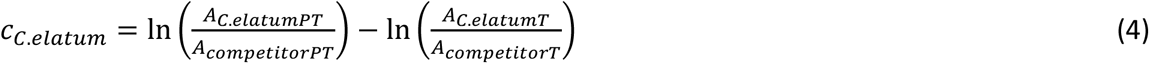

which describes the influence of priming on competition for space: Having experienced a priming cue before the triggering stress, one species might benefit more than the other leading to a shift in the colony ratio compared to the same scenario without priming cue. A value of *c*_*si*_ = 0 refers to no change in colony ratios, i.e. both competitors benefit equally from priming. For a value of *c*_*si*_ > 0, the colony size of *C. elatum* increases more than the one of its competitor. Measuring both, the benefit of *C. elatum* and the competitive shift *c*_*C. elatum*_, allowed us to investigate whether certain parameter combinations affected these values differently, e.g. led to high priming benefit of *C. elatum* but still decreased its competitive strength because the competitor benefitted even more.

## Results

After model fitting, we first validated the model by comparing the simulated output with experimental data of competition treatments not used for model parameterization. We then systematically varied different traits of an artificial species competing with *C. elatum* and assessed the benefit that *C. elatum* gained from priming. Additionally, we measured the effect of priming on competition strength under stress conditions.

### Model Validation

With our model, we could well predict the growth in competition of four of five fungal pairs under stress with and without preceding priming cue (see Fig. 1 and Fig. S1). In these successful cases, the effects of interactions and stress were additive. When competing with *M. elongata*, however, the model underestimated the performance of *C. elatum*: While under control conditions, *M. elongata* overgrew *C. elatum* in the experiment and thus dominated strongly, under stress, *M. elongata* changed its behavior and could no longer overgrow *C. elatum*. This led to a stronger benefit of *C. elatum* instead of the expected additive effects. For the sake of simplicity, however, our model does not yet take into account interactions between competition and stress nor alternative forms of fungal interactions such as overgrowth.

**Figure 1.**
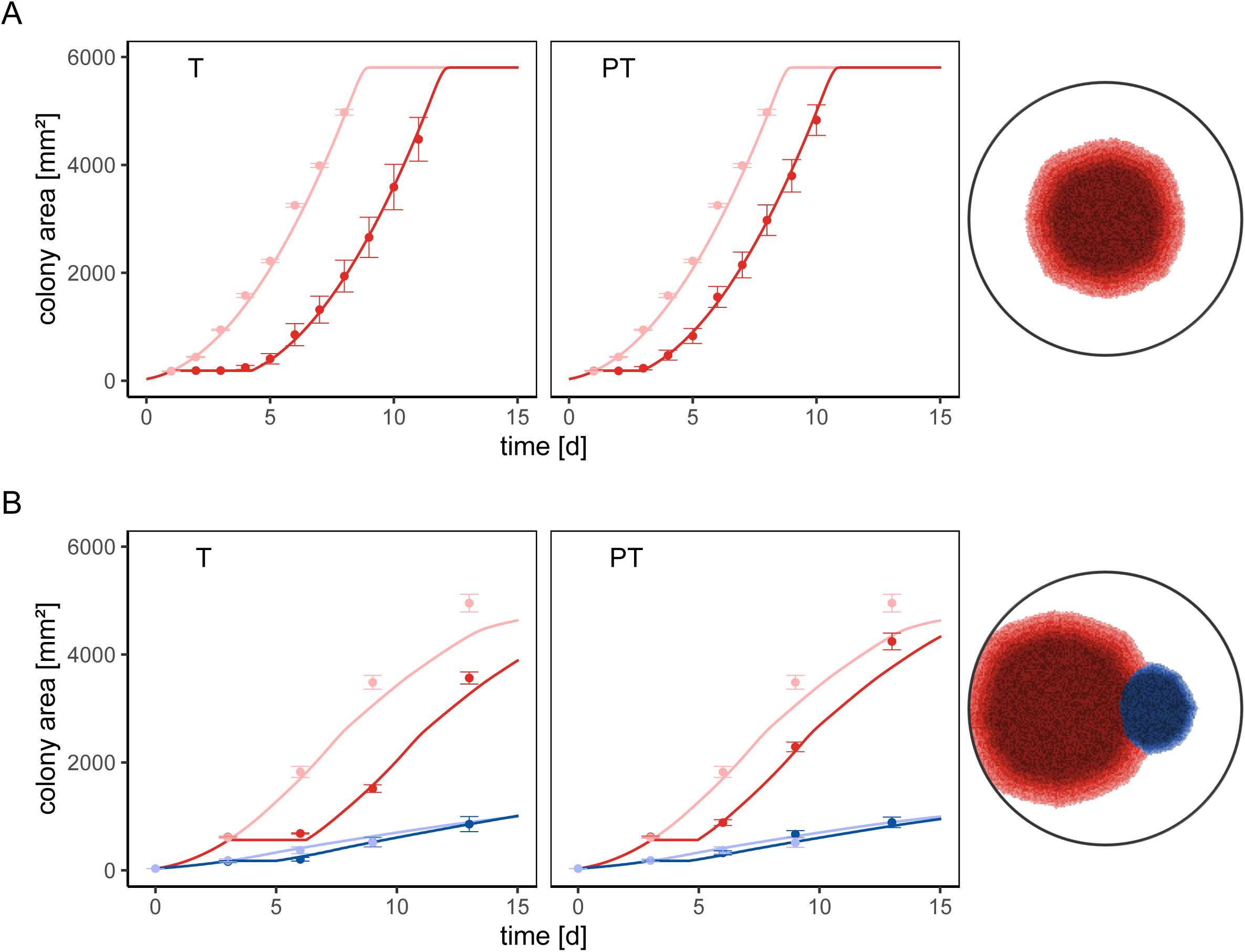
Measured and simulated growth dynamics of *C. elatum* in A) isolation and B) competition with *P. sapidus*. Points describe empirical measurements, and lines are the corresponding simulation model output. Light shades represent the control treatment, while darker shades represent the respective stress treatments (stressed, T, or primed and stressed, PT). Error bars show the standard error of the mean of the observed data. Examples at the right show the corresponding output of the cellular automaton model at day eight.

The priming benefit of *C. elatum* predicted by the simulation model was within the range of variation of the observed benefit for all five pairs (Fig. 2b). By comparing T and PT treatment, the benefit quantifies the effect of priming, canceling out stress effects: The underestimation of the performance of *C. elatum* competing with *M. elongata* thus does not affect the model prediction on priming benefits.

**Figure 2.**
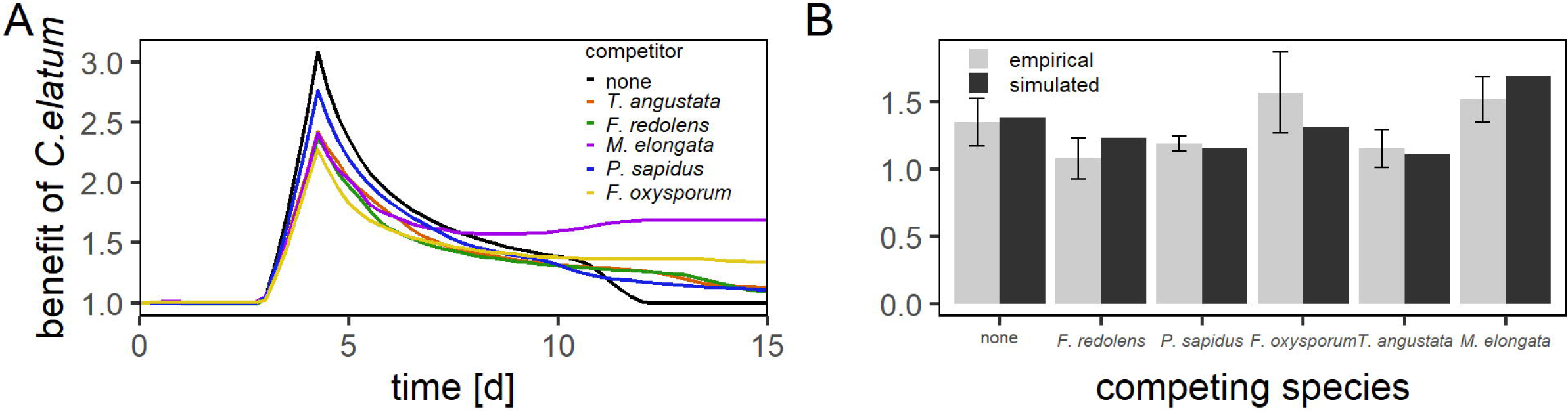
Priming benefit of *C. elatum.* A) Simulation of the priming benefit of *C. elatum* over time in isolation or competition. B) Comparison of the priming benefit of *C. elatum* in isolation or competition with one of five other soil fungi. Values represent the observed benefit at the last day of measurements and the simulated benefit for the same day.

### Priming benefits

When simulating the development of priming benefits over time, a consistent pattern emerged for both isolated and competitive growth (Fig. 2a): The benefit increased just after primed *C. elatum* restarted growth after the lag phase, and reaches a maximum when the non-primed lag phase ended. The subsequent decrease in benefit results from the simultaneous increase each colony’s areas leading to a smaller relative difference between their area.

For isolated growth, the immediate benefit was larger than under competition, because an isolated colony could expand unimpeded and benefit strongly from the shortened lag phase, while under competition, this space might be already occupied by a competitor. A competitor would have already claimed part of the space a species could grow on. The final benefit, however, was lowest in isolation, because without competitors there was no advantage in claiming space earlier, as eventually all available space would be overgrown. This final benefit was largest when *C. elatum* faced one of the two most competitive, i.e. fastest growing, species investigated: *M. elongata*, which exhibits no primeability, followed by *F. oxysporum*.

When we systematically varied fungal traits, for all combinations of traits within the investigated parameter space, priming was beneficial (i.e. benefit > 1, Fig. 3) eight days after inoculation, since priming involved no costs. However, under competition with a highly primeable and stress-susceptible competitor, priming was only marginally beneficial, especially when the competing species was fast-growing. Conversely, we observed the highest benefit when *C. elatum* faced a stress-resistant and only moderately primeable competitor. Here, the negative effect of fast growing competitors was reversed (upper right vs. lower left panel of Fig. 3). A very susceptible competitor with high primeability strongly reduced its lag phase under priming. The faster that opponent grows, the more *C. elatum* will suffer from its gain in growing time. If, however, the opponent is not very primeable, a priming cue will be of no great advantage to that species. In this case, *C. elatum* will benefit even more if the competitor is fast-growing.

**Figure 3.**
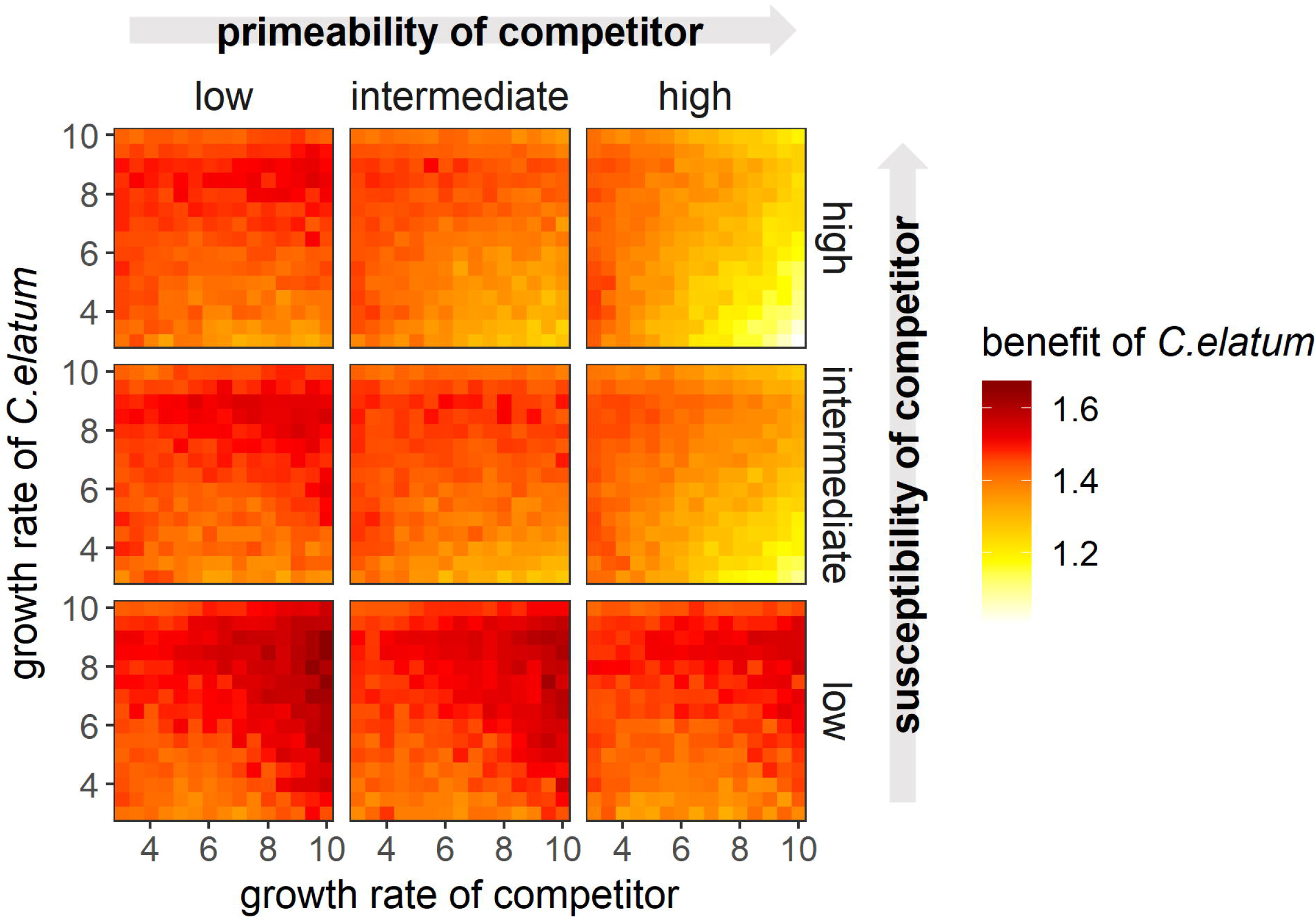
Priming benefit of *C. elatum* in competition with an artificial species. Benefits are shown for different trait combinations eight days after stress treatment. Levels of susceptibility correspond to different lengths of a stress-induced lag phase: low = 0.5 days, intermediate = 1.5 days, high = 2 days, and levels of primeability correspond to the reduction of this lag phase under priming conditions: low = 25%, intermediate = 50%, high= 100%.

Fifteen days after inoculation, when the Petri dish was filled and a steady state was reached, priming was not beneficial (i.e. benefit < 1) in case of a fast growing, highly primeable competitor with intermediate or high stress susceptibility (Fig. S2). During phases of growth, space that is lost to a more primeable competitor can still be compensated by colonizing empty space elsewhere. In this case, it can be more beneficial not to be primed at all, if space is limited.

### The effect of priming on competition

Analogous to the investigation of priming benefits, we measured how the colony ratio between *C. elatum* and its competitor changed depending on different fungal traits (Fig. 4 and Fig. S3). A positive value indicates that priming favors *C. elatum* more than its competitor. Again, the least favorable conditions arose when *C. elatum* faced stress-susceptible but primeable opponents. However, because of its intermediate primeability, for most investigated trait combinations, *C. elatum* benefited stronger than its competitor, as the long stress-induced lag phase of *C. elatum* is reduced substantially.

**Figure 4.**
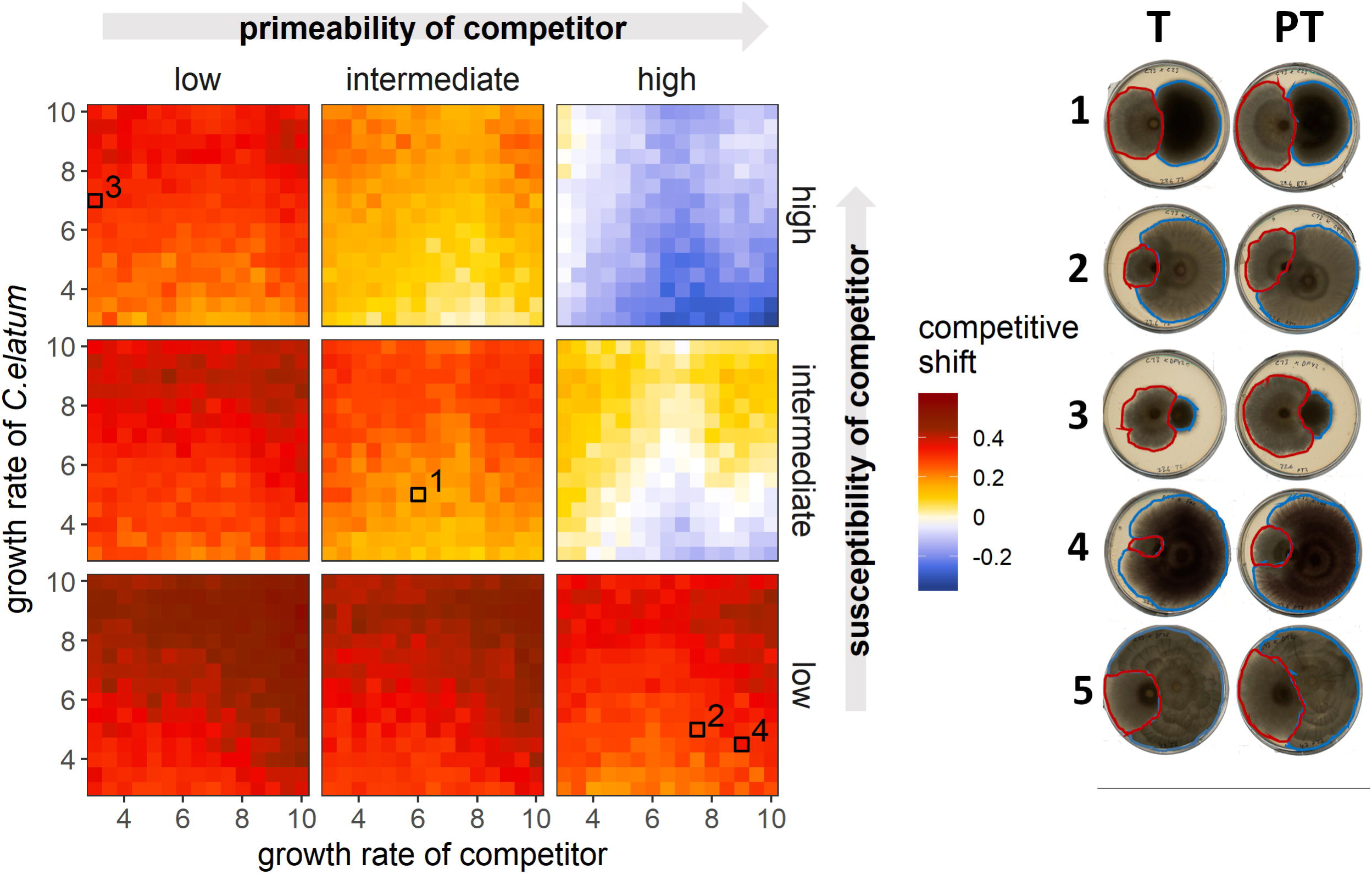
Competitive shift of *C. elatum* in competition with an artificial species. The shifts in competition are shown eight days after the stress treatment. Red shades indicate a shift in favor of *C. elatum*, and blue shades a shift favoring its competitor. Photos show exemplary pairwise cultures grown in the laboratory: Each pair is assigned the respective shift of competition predicted by the simulation model according to the parameter values of the competitor and growth of *C. elatum*. The pairs shown are *C. elatum* (red) competing against (blue) 1. *F. redolens,* 2. *T. angustata,* 3. *P. sapidus* 4. *F. oxysporum,* 5. *M. elongata. M. elongata* is fast growing and not primeable, and is not represented in the visualized parameter space. Levels of susceptibility correspond to different lengths of a stress-induced lag phase: low = 0.5 days, intermediate = 1.5 days, high = 2 days, and levels of primeability correspond to the reduction of this lag phase under priming conditions: low = 25%, intermediate = 50%, high= 100%.

When comparing figure 3 and 4, it is notable that while in many cases priming conferred a moderate benefit to *C. elatum*, the competitive shift in these cases was variable: facing a primeable but stress susceptible and slow competitor, *C. elatum* moderately benefitted from priming. However, since the competitor’s benefit is higher, the colony ratio was shifted in favor of the competitor. If on the other hand the competitor was less primeable, the absolute benefit of *C. elatum* was similar as before, but this time leading to a positive shift in the colony ratios in favor of *C. elatum*.

## Discussion

We successfully developed a cellular automaton model that reproduces growth of competing fungi in a Petri dish under priming and heat stress conditions. With this model, we assessed how different fungal traits such as stress susceptibility and primeability influence the species-specific benefit and competition outcomes.

### The priming response of fungi

For all experimentally treated species, a heat-induced no-growth phase was observed in the experimental data, and for four of six species, the post-lag growth phase was not significantly different from the control growth. Priming did not affect the growth of any of the species, but instead reduced the duration of the phase without growth.

An analytical study by Wesener and Tietjen (2019) using coupled differential equations of microbial growth showed that stress of short duration is best met with an early defense and that a primed stress strategy is most successful when further shortening the response. The current study confirms this pattern, as the fungi were treated with two-hour pulses of heat instead of prolonged periods of warming, and the primed colonies restarted growth earlier than those that had not been primed. Especially for species with a regeneration phase less than a day (*F. oxysporum* and *F. redolens*) the temporal resolution of measurements after stress should be increased to enable differentiation between an immediate but slow reversal to control level growth or a lag phase with no growth and “switch-like” change. Additionally, longer durations of heat stress could be applied to test whether the fungal stress response types differ for different types of heat stress.

In all cases, heat stress affected growth. Additionally, in some species combinations it could also qualitatively alter the type of interaction between competitors, such as changing overgrowth to deadlock. This is in line with previous findings (Hiscox, Clarkson, *et al.*, 2016) and could be a valuable extension to our simulation model.

### Priming costs

In this study, we aimed at accurately imitating growth dynamics of fungi under priming conditions. We ended at not implementing any costs of priming, as there was little evidence under laboratory conditions that costs of priming are realized as reduced growth. Because priming usually involves the transient production of precursor molecules or transcription factors rather than the accumulation of resistance compounds, priming costs are generally expected to be low (Heil, 2014) and might be hard to quantify. Especially under laboratory conditions, costs of induced resistance can be overseen, e.g. when they manifest as ecological costs (Heil, 2002).

The distribution of resources between growth, resistance and reproduction is central to ecological theory, and any defense strategy must entail some costs (Harvell, 1990; Schulenburg *et al.*, 2009; Crowther *et al.*, 2014). A priming mechanism without costs would not bear any risks, and even in environments with low stress predictability (leading to organisms reacting to a priming cue, which is not followed by a triggering stress), priming would be of no disadvantage and would be ubiquitous in nature. To our knowledge, there is no study that investigated the costs of priming in microbes. Studies on priming costs in plants differed in their results for different species and priming cues, finding no direct costs of priming (Perazzolli *et al.*, 2011), costs realized as growth reduction (Hulten *et al.*, 2006), or reduced rhizome production (Yip *et al.*, 2019). Priming costs in fungi might thus also not be manifested in reduced growth, but rather in reduced spore production or competitive strength. Therefore, we want to stress the need of research on costs of induced resistance in microbes, which is necessary to fully comprehend the benefits and potential trade-offs of priming.

### The benefit of priming

Because we did not implement any priming costs, during the growth phase priming is generally beneficial for *C. elatum* in all investigated scenarios. Therefore, we focus rather on the magnitude and not on the presence of this benefit.

Our results show that the benefit of priming under competition is highly dependent on fungal traits such as primeability, stress susceptibility and growth, as well as the time point during community buildup. We could show that depending on these factors, priming might not always more beneficial under competition compared to the isolated benefit. Even when priming itself is of low direct benefit for a given species, it can still shift dominance in favor of that particular species, depending on the traits of its competitors. Priming can influence competition between species by shifting dominance towards one of the competitors, especially favoring primeable species that otherwise show a competitive disadvantage (i.e. not stress-resistant or slow-growing). Priming can thus be beneficial when taking into account the change of the community structure and the resulting fitness of competing species, making it difficult to infer priming effects in a community from effects measured on species in isolation.

*C. elatum* exhibits moderate primeability and shows the longest stress-induced lag phase of the six investigated species. As a result, priming has the potential to strongly shorten its lag phase and thus to be highly beneficial in comparison to its competitors with lower susceptibility or lower primeability. Because the competition for space in fungal communities is effectively competition for gaining access to nutrients (Boddy, 2000), it is of particular importance when recolonizing new territory. A primed stress response that allows an organism to occupy empty space earlier than its competitors therefore leads to the additional advantage of claiming space that would otherwise be colonized by another species. The order of species arrival in community assembly affects community structure and function (Fukami, 2015) and priority effects have been shown to be a common influence on fungal communities (Kennedy *et al.*, 2009). These priority effects can even be of increased importance when species further change environmental conditions or resource availability for later species via niche modification (Fukami, 2015). Environmental factors such as temperature have been shown to influence assembly of fungal community members (Hiscox *et al.*, 2015; Hiscox, Clarkson, *et al.*, 2016; Hiscox, Savoury, *et al.*, 2016). Therefore, heat priming can potentially influence the order of community assembly by letting certain species grow earlier than others.

Priming might not only affect community composition via community assembly, but also directly influence community structure: Sensitivity of microbial communities to disturbances is common, as they rarely return to pre-disturbance composition and reach alternative stable states (Shade et al. 2012; Schimel, Balser, and M. Wallenstein 2007; Allison and Martiny 2008). Environments with fluctuating temperature increase species number in fungal communities (Toljander *et al.*, 2006), and post-stress communities can transiently consist of species that are generally more resistant to stress because of their higher likelihood to survive past stress events (Evans and Wallenstein, 2012; Jurburg *et al.*, 2017). Priming, however, can influence community resistance, if less resistant but instead primeable species persist in a community. Stress responses at an individual level, such as priming, might therefore interact with legacy effects arising from pre-disturbance community composition (Meisner *et al.*, 2018), resulting in communities with different functions or stress resistance.

With this study, we could show that the effect of priming on a species and on the community structure is not consistent, but highly varies in strength depending on the primeability and stress susceptibility of other members in the community. It is therefore essential to understand how the community context and different species traits affect priming to comprehend its influence on community assembly. We therefore want to stimulate further research on inducible stress defenses in microbes and their effects on community development, as well as the effect of the community context on stress defense mechanisms.

## Acknowledgements

We are grateful to the German Research Foundation (DFG) for funding our Collaborative Research Centre 973 ‘Priming and Memory of Organismic Responses to Stress’ (www.sfb973.de).

