## Supplementary Material for "Stress priming affects fungal competition – evidence from a combined experimental and modeling study"

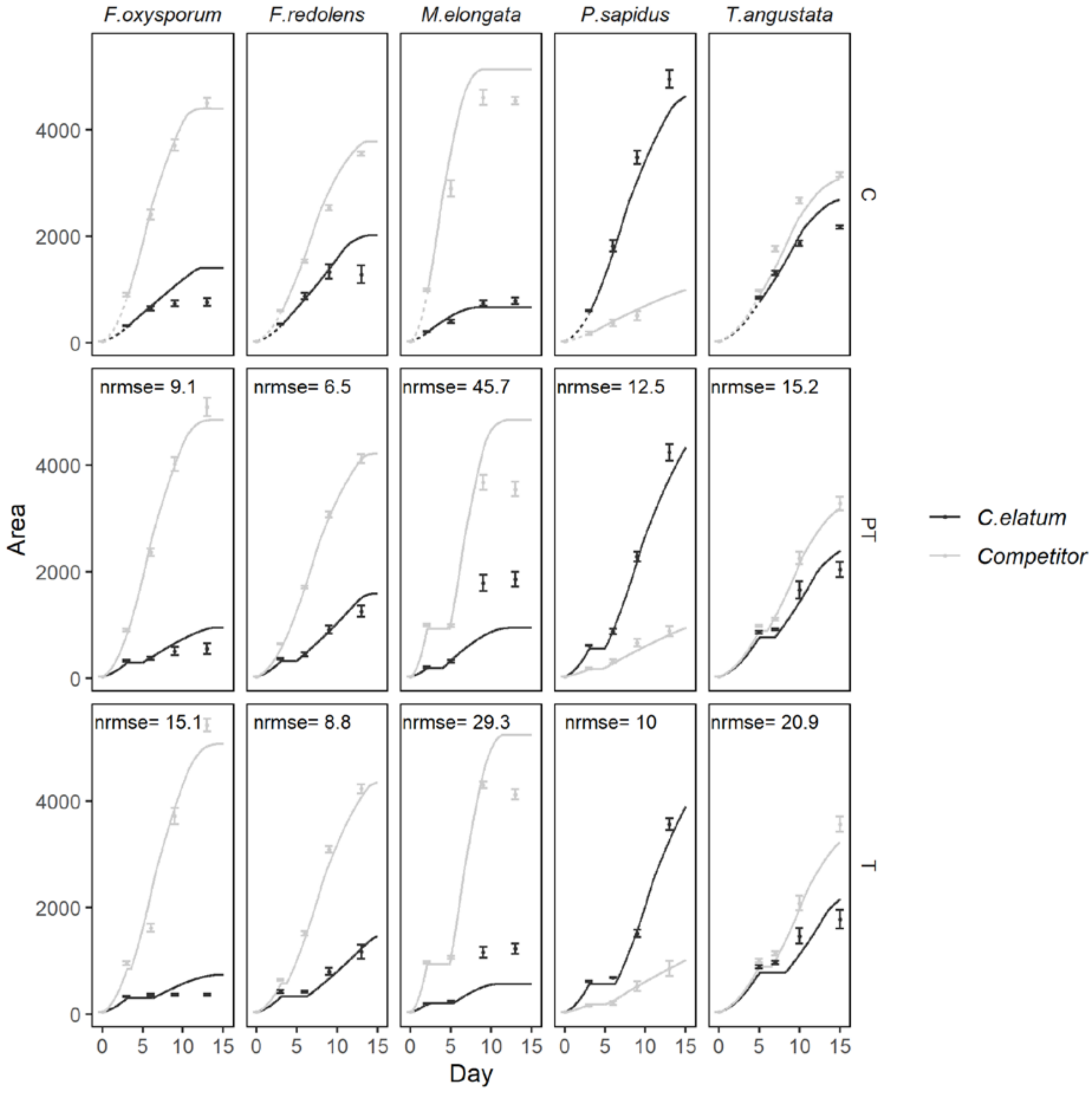


**Figure S1** Growth dynamics of *C. elatum* competing with five other soil fungi. Points describe empirical measurements, and lines are the corresponding simulation model output. C = control treatment, T = stressed treatment, PT = primed and stressed treatment. Error bars show the standard error of the mean of the observed data. Nrmse is the normalized root mean square error of the simulation model for the respective pair.

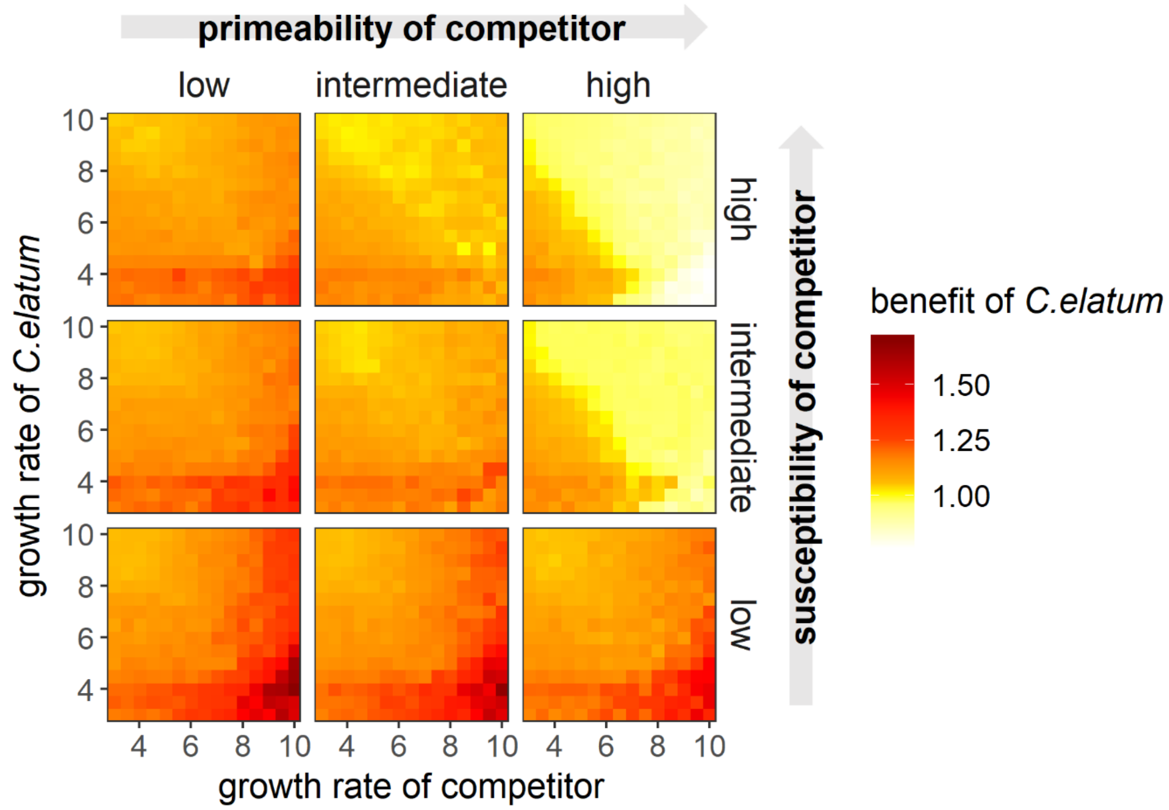


**Figure S2** Priming benefit of *C. elatum* in competition with an artificial species. Benefits are shown for different trait combinations fifteen days after stress treatment at the end of the simulation. Levels of susceptibility correspond to different lengths of a stress-induced lag phase: low = 0.5 days, intermediate = 1.5 days, high = 2 days, and levels of primeability correspond to the reduction of this lag phase under priming conditions: low = 25%, intermediate = 50%, high= 100%.

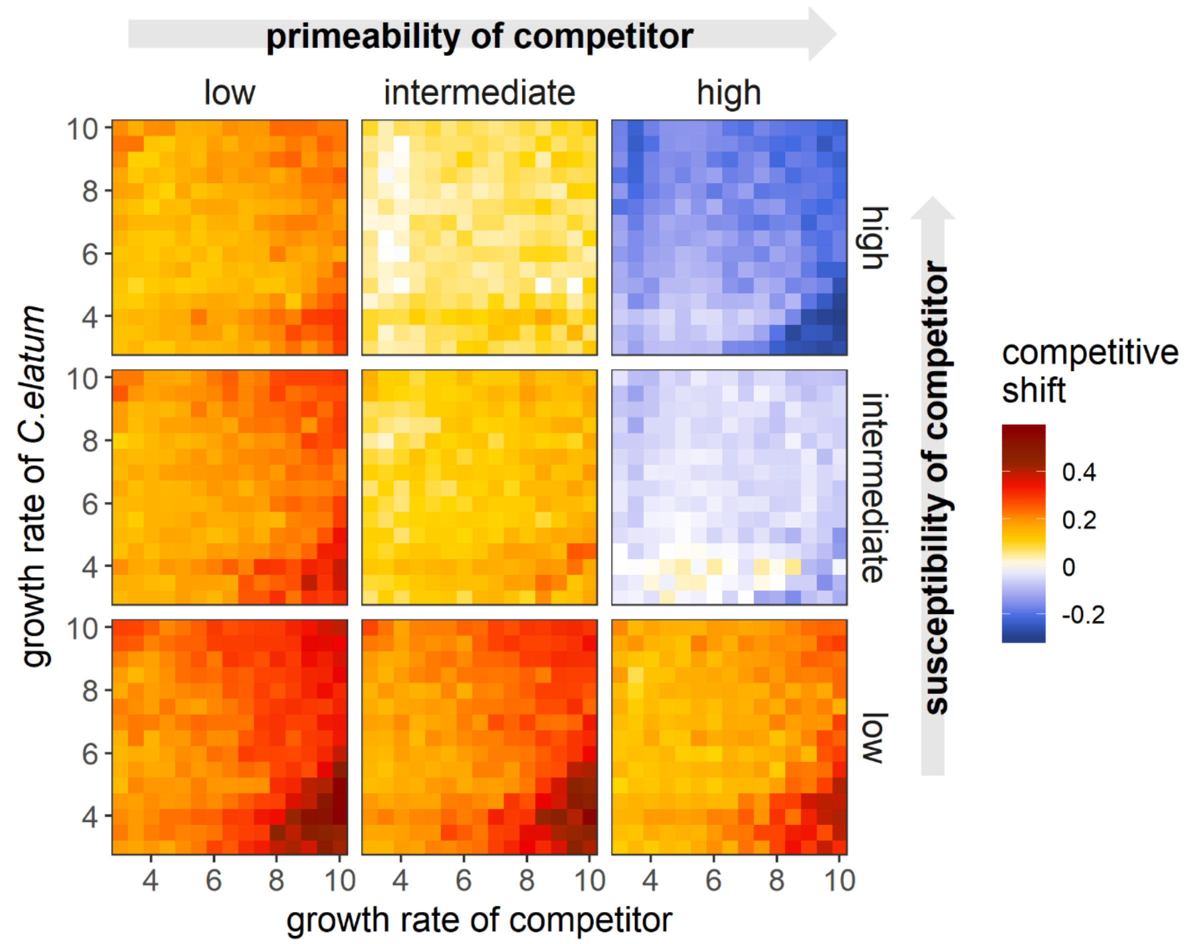


**Figure S3** Competitive shift of *C. elatum* in competition with an artificial species. The shifts in competition are shown fifteen days after the stress treatment. Red shades indicate a shift in favor of *C. elatum*, and blue shades a shift favoring its competitor. Levels of susceptibility correspond to different lengths of a stress-induced lag phase: low = 0.5 days, intermediate = 1.5 days, high = 2 days, and levels of primeability correspond to the reduction of this lag phase under priming conditions: low = 25%, intermediate = 50%, high= 100%.
